# Defending the Undefendable: Male Territorial Behaviour and Mating System in Monogamous Primates

**DOI:** 10.1101/354118

**Authors:** R.I.M. Dunbar

**Affiliations:** Department of Experimental Psychology University of Oxford South Parks Rd Oxford OX1 3UD, UK email

**Keywords:** monogamous primates, female dispersion, territory size, male reproductive strategies

## Abstract

It has been suggested that monogamy evolves when females forage alone and are overdispersed, such that males cannot defend more than one female at a time. I test the underlying assumption that the females of monogamous anthropoid primates are overdispersed in three different ways, and compare the results with data for several polygynous primate genera. First, I show that monogamous primates do not have per capita territories that are significantly larger than those of polygynous taxa. Second, given their day journey length and the Mitani-Rodman equation (Mitani & Rodman 1979), males of most monogamous species could easily defend areas large enough to allow them to monopolise 5-6 females. Finally, I use a model of male mate searching strategies to show that, unlike the males of polygynous species, the males of monogamous species would sire more offspring by adopting a roving male form of polygyny when females are dispersed. The opportunity cost that monogamous males incur is typically more than five times the reproductive success they have by being obligately monogamous, suggesting that the selection pressure preventing them from pursuing a roving male strategy is very considerable. Given that biparental care always follows the adoption of monogamy in primate evolution, the only viable explanation for monogamy would seem to be either high predation risk or high infanticide risk.

## Introduction

The evolutionary forces that select for monogamy in mammals have continued to attract debate. Generally speaking, three principal hypotheses have been proposed, namely (1) biparental care, (2) female (over-)dispersion and (3) infanticide risk (Opie et al. 2013). The role of female dispersion has strong comparative support as the likely explanation for mammals in general (Komers & Brotherton 1997; Brotherton & Komers 2003; Lukas & Clutton-Brock 2013). Females are forced to disperse because they need large ranges in which to find sufficient food resources to maintain themselves and their offspring. Many monogamous mammals (especially among the ungulates) are small-bodied, specialist feeders dependent on patchy high quality resources, and the likelihood that females are over¬dispersed is very plausible in their case (though, it may be noted, no evidence has ever been adduced to show that this is in fact so). However, other hypotheses, notably protection against infanticide, have been favoured in the case of primates (e.g. van Schaik & Dunbar 1990; Opie et al. 2013), although the female dispersion hypothesis has continued to be advocated (e.g. Lukas & Clutton-Brook 2013).

The plausibility of the female dispersion hypothesis rests on the claim that males cannot defend a territory large enough to encompass the range of more than one female in those species where females range alone (i.e. not in groups). Crucial support would therefore be given to this hypothesis if it could be shown that: (1) the females of monogamous species occupy larger *per capita* territories than females of polygamous species, (2) the males of monogamous species really are unable to defend territories larger than those they actually occupy, and (3) given that their females forage alone, the males of monogamous species would not do better by opting for a roving male mating strategy. I test all three of these assumptions using comparative data for monogamous species of primates.

To test the second assumption, I exploit a relationship first noted by Mitani & Rodman (1979) who showed that primate species are only territorial when the ratio of their day journey length to the diameter of their range area is greater than unity (in effect, they have to be able to traverse their territory with a minimum frequency that allows them to detect and expel intruders). We can use this relationship, following Dunbar (1988, 1995a) to determine the largest area that a male could defend, and then determine how many females would fit into this area given the actual range size that females and their offspring typically occupy. This extension of the Mitani-Rodman equation is a simple logical extrapolation of their formula and rests on the empirical fact that this relationship is a reliable predictor of territorial behaviour. The Mitani-Rodman principle was confirmed by Lowen & Dunbar (1994) using an alternative approach based on the Maxwell-Boltzmann gas dynamics equations. They derived an alternative index that broadly confirmed the Mitani-Rodman relationship while significantly improving its predictability.

To test the third hypothesis, I use a model of male mating strategies developed for apes by Dunbar (2000, 2001). This model calculates the payoff (in terms of the number of offspring sired over the average female reproductive cycle) by a male that pursues a roving male strategy (i.e. searches promiscuously for groups of females to mate) compared to that obtained by a social male who stays permanently with one group of females. A social male can guarantee siring one offspring with every female in his group in this time interval (i.e. one offspring in the case of monogamous males, where female group size is one), while a roving male’s payoff is determined by the likelihood that he will chance upon an unfertilised female in any of the groups he encounters while searching randomly. The payoff for a promiscuous roving male will depend on the male’s search rate (the path he carves out each day, and the number of days for which he does this), the density and size of female groups, and the females’ characteristic reproductive parameters. This model is extremely robust for great apes: it predicts with almost no error variance the proportion of males that are social (i.e. are found with groups of females on any given occasion) in populations of chimpanzees, gorillas and orang utans. More importantly, the regression line for the data passes through the point of equivalence where males should be undecided whether or not to be social because the payoffs are equal (Dunbar 2000, Fig. 1).

For comparison, I include data for two genera of territorial polygynous primates that live in unimale groups (*Colobus* and *Cercopithecus*) and one non-territorial polygynous genus (*Gorilla*). Although multimale groups do occur in all three of these genera, the typical social group is unimale (Fashing 2007; Enstam & Isbell 2007; Robbins 2007). *Cercopithecus* also includes two species (*C. cephus* and *C. neglectus*) that appear to be monogamous (they have unusually small groups for a cercopithecine monkey, typically one male and one female), and I analyse their data separately. As a further comparison, I also include 11 species of ‘solitary’ nocturnal prosimians (two lemurines, three lorisids and six galagines) whose mating system is essentially roving male promiscuity: males defend large territories that enclose the territories of several females who forage alone. These comparator taxa provide us with an important reality check on the models because their indices should favour group-living polygyny rather than roving male polygyny for the three catarrhine genera and roving male polygyny rather than monogamy for the prosimians. If the models cannot correctly predict the behaviour of these taxa, they are unlikely to do so reliably for the monogamous species.

## Methods

Data on day journey length, territory size, social group size, mean number of females per group and interbirth interval were sourced from Campbell et al. (2007), the most comprehensive and up to date compilation available, for 14 monogamous anthropoid genera (hylobatids, callitrichids, the small-bodied cebids, and two species of the genus *Cercopithecus*), three genera of comparator anthropoids (*Colobus*, *Cercopithecus* and *Gorilla*) and 11 species of promiscuous/solitary nocturnal prosimians. The species included in the sample are listed in the Appendix. Altogether, 55 species from 16 genera are represented in the main sample. Campbell et al. (2007) do not provide data on *Gorilla*, so these are taken from Doran & McNeilage (1998), references cited therein and a special issue on lowland gorillas (Doran-Sheehy & Boesch 2004). For 11 species of ‘solitary’ nocturnal prosimians, I obtained data on average male territory size and average female territory size from Campbell et al. (2007). Data on group density are from Campbell et al. (2007) in the first instance, and Smuts et al. (1987) for taxa for which Campbell et al. (2007) do not provide data. Data on interbirth intervals are used only if they derived from wild populations; where possible, the value given for each specific study site is used, otherwise I used the species mean value. Most species are represented by 1-3 populations, with just three species having more populations sampled (*Colobus guereza*: 5; *Hylobates lar*. 4; *Symphalangus syndactylus*: 4).

The callitrichids are somewhat anomalous because, although they have traditionally been considered to be monogamous, their mating systems is in fact more complex and variable, with individual groups being able to switch between monogamy, polygamy, polyandry and polygynandry (Goldizen 1988; Dunbar 1995a,b; Digby et al. 2007; Opie et al. 2013). Nonetheless, for present purposes, I include them in the monogamous sample because they normally have only a single breeding pair, irrespective of the composition of the social group.

Mitani & Rodman (1979) showed that a simple geometric index based on day journey length and territory area could be used to differentiate reliably between territorial and non-territorial primate species. Their index is the ratio of day journey length to the diameter of the range area (assuming this to be a perfect circle): species are only territorial when this ratio is greater than unity. (Although not all species with indices greater than 1 are territorial, species with indices smaller than 1 are never territorial.) In effect, so long as the group can, in principle, traverse from one side of its range to the other easily during a day’s foraging, it is able to defend its range area as a territory. Lowen & Dunbar (1994) developed a more complex based on the Maxwell-Boltzman gas dynamics equation: this estimates the number of times that a group would ‘hit’ its range boundary during a day when ranging randomly. Although it has a significantly higher goodness of fit than the Mitani-Rodman equation (and thus provides further confirmation of the principle), it does not provide such an intuitively simple criterion for defining the maximum defendable area. I therefore use the Mitani-Rodman relationship for the analyses in this paper, but rely on the fact that it has independent confirmation to justify doing so.

Mitani & Rodman (1979) showed that a species is territorial so long as:

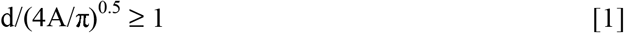

where *d* is the day journey length and *A* the home range area. We can use this inequality to determine *A*_*max*_, the maximum size of territory that a male can defend given its typical day journey length, by inverting inequality [1] and setting it equal to 1:

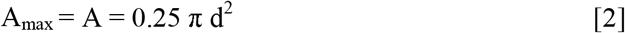

Following van Schaik & Dunbar (1990) and Dunbar (1995a), we can use Eq. [2] to calculate the number of reproductive females a male could expect to include within its maximally defendable territory by dividing *A*_*max*_ by the size of territory required to support one female and her offspring. The area that one female needs to support herself and her offspring can be estimated as 67% of the territory that the species has at a given location (allowing the rest to support the male). The assumption that a female and her offspring need 67% of the territory to provide for their foraging needs is based on the observation that monogamous groups typically consist of an adult male, an adult female and two immatures (who are treated as equivalent to half the adult body mass each). The number of females a male at a particular location can expect to monopolise is thus:

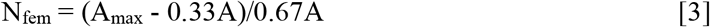

where *N*_*fem*_ is the number of females a male can expect to monopolise in a territory of size *A*_*max*_ and *A* is the observed territory size at that site.

To calculate the number of defendable females that males in the polygamous species could have, I calculated *A*_*max*_ as in Eq. [2], and then calculated the defendable number of females by determining the proportion of the group territory a single female and her offspring require as:

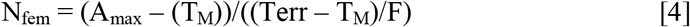

where *F* is the observed mean number of females in a group and *T*_*M*_ is the proportion of the group territory required for the male, namely:

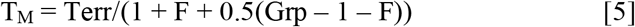

As with Eq. [3], Eq. [5] allocates the equivalent of half the adult body mass to each immature animal. In effect, I partition the group’s observed territory between the adults in the group (one male and *F* females, with their associated offspring), subtract the male’s share from the defendable area and then divide the remainder by 1/*F*(i.e. one female’s share).

The nocturnal prosimians operate a form of roving male polygyny, and I therefore assume that the male’s observed territory size represents the largest area he can defend. To determine how many females a male can monopolise, I first calculated the number of females the male could expect to have within this territory by dividing the male territory by the female territory, and then apportioning the male’s territory between this number of females, an equal number of offspring adult-equivalents and the male. I removed the male’s share of this territory from his total territory size and then divided the resulting value by the observed female territory size.

Following Dunbar (2000), the number of offspring that a roving male can expect to sire during the average female reproductive cycle, *E(f)*, is given by:

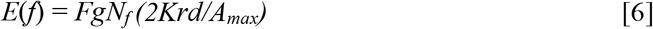

This is made up of two components: the number of times that a roving male can expect to encounter groups of females, *N*_*f*_*(2Krd/A_max_)*, and the expected number of fertilisable females per group, *Fg*. The first component consists of two parts: the probability that a randomly searching male will encounter a female group during an average reproductive cycle, 2*Krd* (calculated as the proportion of the total territory, *A*_*max*_, that he can defend, given a daily search path of *d* km length [from Eq. 2], a detection distance either side of his path of travel of *d* km [taken to be 200m] and a reproductive cycle [or interbirth interval] of length *K* days) and the number of female groups in his maximally defendable territory (*N*_*f*_). The relationship does not specifically depend on Eq.[2] and the Mitani-Rodman relationship; it depends only on the density of female groups and the proportion of land surface that a male can search each day when foraging randomly; however, it is convenient to define that area as *A_max_* and use that to delimit the area over which female groups might be encountered. The mean number of fertilisable females in a typical female group is the expectation (or mean) of a Poisson process with parameter *g* (the probability that a female will be at risk of conception on any given day) and the mean number of females in a female group (*F*). The model assumes that a female is receptive (i.e. fertilisable and not merely in oestrus as conventionally defined) for about 5 days on each of three menstrual cycles during any given reproductive cycle (as is typical for primates), hence *g*=15/*K* The model uses the reproductive cycle length (or interbirth interval) as its time base because a male who is social (i.e. stays permanently with a group of females) can expect to sire exactly one offspring with each female during this interval, thus providing a very convenient baseline for comparison.

## Results

### Comparative Territory Sizes

The distribution of territory sizes for the monogamous genera are plotted in Fig. 1. Territory sizes per adult female for colobine and cercopithecine genera that live in unimale groups are shown for comparison. In both cases, I have removed that share of the group territory that can be ascribed to the male, following Eq.[5]. I have not included the data for *Gorilla* since these are likely to be confounded by the enormous body size difference in their case. Although there is a significant difference in mean territory size per female among the various genera (*F*_11,56_=10.49, p<0.001), in fact this is entirely due to *Leontopithecus* which differs significantly from all other genera (Scheffé *post hoc* tests: p≤0.008) except *Callimico* (p=0.293); none of the other pairwise comparisons are significant (Scheffé *post hoc* tests: p≥0.920). In other words, although the females of monogamous species may sometimes have slightly larger *per capita* territories than those of polygynous species (as Fig. 1 suggests), their territories are not consistently larger. It thus seems that monogamous females are not overdispersed, even if they typically forage separately from other adult females.

**Fig. 1.**
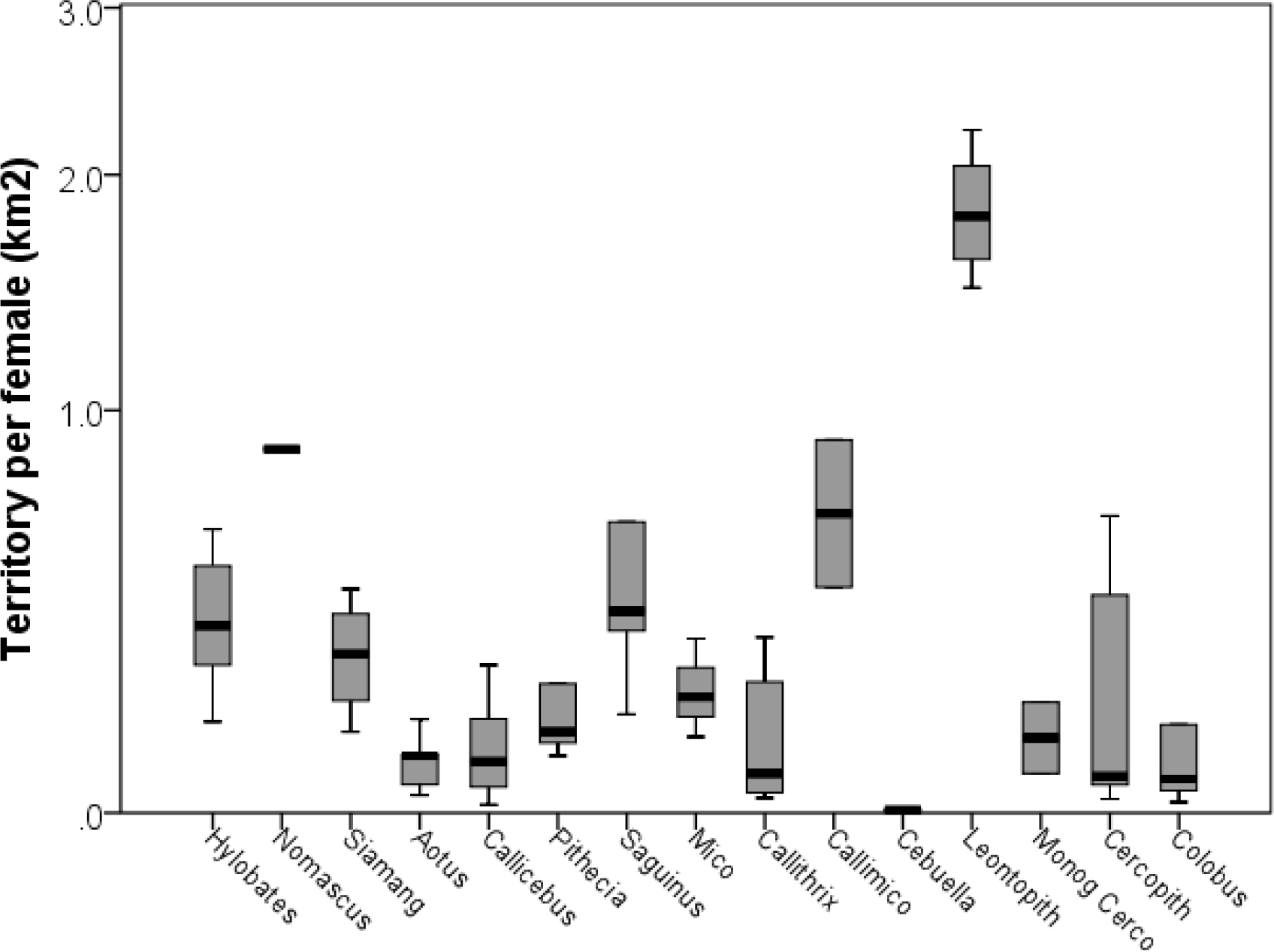
Median (±50% and 95% ranges) observed territory size (km^2^) per adult female group member for all monogamous genera and the two comparator polygynous genera (*Cercopithecus* and *Colobus*). Monog Cerco refers to the two putatively monogamous *Cercopithecus* species (*C. cephus* and *C. neglectus*). Only *Leontopithecus* differs significantly from the other genera (except *Callimico*).

### Maximally Defendable Territory Size

Fig. 2 plots the predicted number of females that a territorial male from each of the habitually monogamous primate genera could expect to have within its maximally defendable territory. The equivalent values for polygamous genera that live in unimale groups (colobines, cercopithecines and gorillas) and for nocturnal Prosimians that live in dispersed social systems are plotted for comparison. It is clear that males of all of the habitually monogamous genera would easily be able to defend a territory large enough to encompass the ranges of several females. Most would typically be able to monopolise around 5-6 breeding females. For none of the three polygynous genera would males gain access to significantly more females than they actually do (matched sample *t*-tests comparing number of monopolisable females against actual female group size: *Cercopithecus*, p=0.058; *Colobus*, p=0.477; *Gorilla*, p=0.993), although cercopiths clearly do better in this respect than the others. (*Cercopithecus* groups are significantly larger than both *Colobus* and *Gorilla* groups and their males are often unable to prevent invasions by groups of males [Henzi & Lawes 1987]; they seem to be on the cusp of what is defendable between unimale and multimale groups [Andelman 1987]). By contrast, none of the nocturnal Prosimian males would benefit by being social (in effect, monogamous): on average, all do better by pursuing a roving male strategy (p=0.022), as in fact they do.

**Fig. 2.**
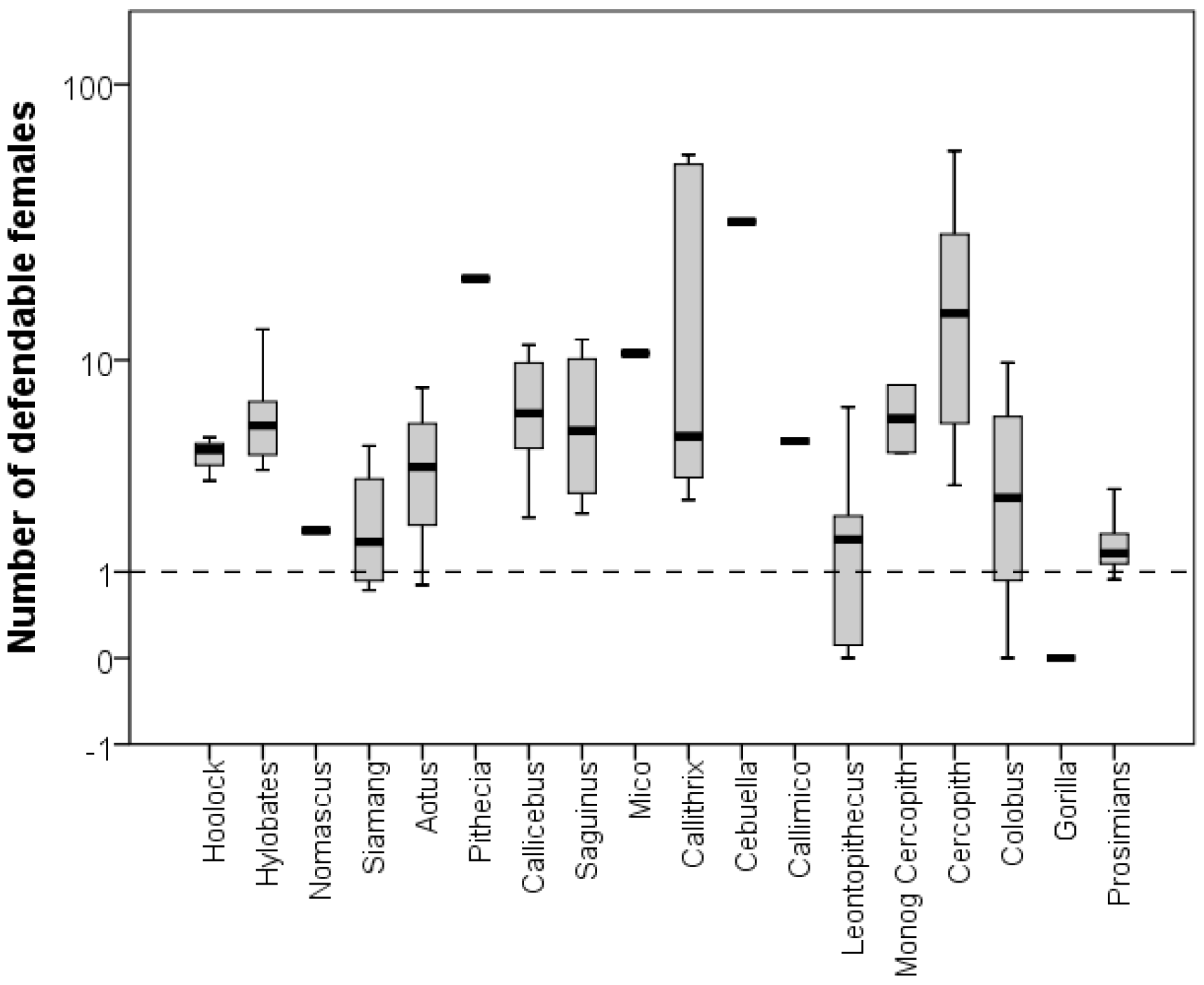
Median (±50% and 95% ranges) of the number of defendable females for individual monogamous genera, the three comparator polygynous genera (*Cercopithecus*, *Colobus* and *Gorilla*), and the promiscuous nocturnal prosimians, as predicted by the Mitani-Rodman equation. The dashed line indicates the benchmark of monogamous taxa (one female). For the polygynous genera, the median observed number of adult females per group is 7 for *Cercopitjhecus*, 3.8 for *Colobus*, and 4 for *Gorilla*.

In calculating the number of females that could be fitted within a maximally defendable territory, I assumed that monogamous groups typically consisted of four individuals (one adult male, one adult female and two offspring whose body mass was half that of an adult on average). This assumption does not always hold for callitrichids, who typically twin and, at least in some cases, can have up to two litters a year who remain in their natal group to act as helpers-at-the-nest (Digby et al. 2007). Eq. [3] will therefore tend to underestimate the share of the territory required by the female and may result in the number of females that a male could have in a maximally defendable territory being overestimated. *Callithrix* groups average around 8.5 members, with typically a single breeding pair; together with the offspring of various ages, this would give an effective metabolic group size of around 6.5 adult-equivalents. Increasing the proportion of the territory that a breeding female requires for herself and her offspring to 85% reduces the number of females a *Callithrix* male would have access to by, on average, just 17% and does not change the general result in Fig. 2. It would take a very much more substantial increase in group size to wipe out the advantage to a polygynous male, and that would be achievable only by having multifemale groups (in other words, a polygynous mating system).

A second possible confound is that Eq.[4] assumes that group territories are densely packed. If groups were overdispersed, and their density was less than one-fifth of that assumed by continuous packing of territories, then polygyny would not be a viable strategy. Fig. 3 plots the mean density of groups for the monogamous taxa, based on actual field estimates. The median observed group density for monogamous species (indicated by the dashed line) is 4.9 groups per km^2^; the median estimated density based on Eq.[4] is 3.3 groups per km^2^ (indicated by the dotted line). With the possible exception of *Cebuella*, none of the densities used in determining the number of defendable females are higher than the observed density; if anything, they are actually lower. This is mainly due to the fact that there is often considerable territory overlap between neighbouring groups in many of these species (Campbell et al. 2007). This makes it unlikely that Fig. 2 overestimates the payoff to promiscuous males; in anything, it probably underestimates the true value.

**Fig. 3.**
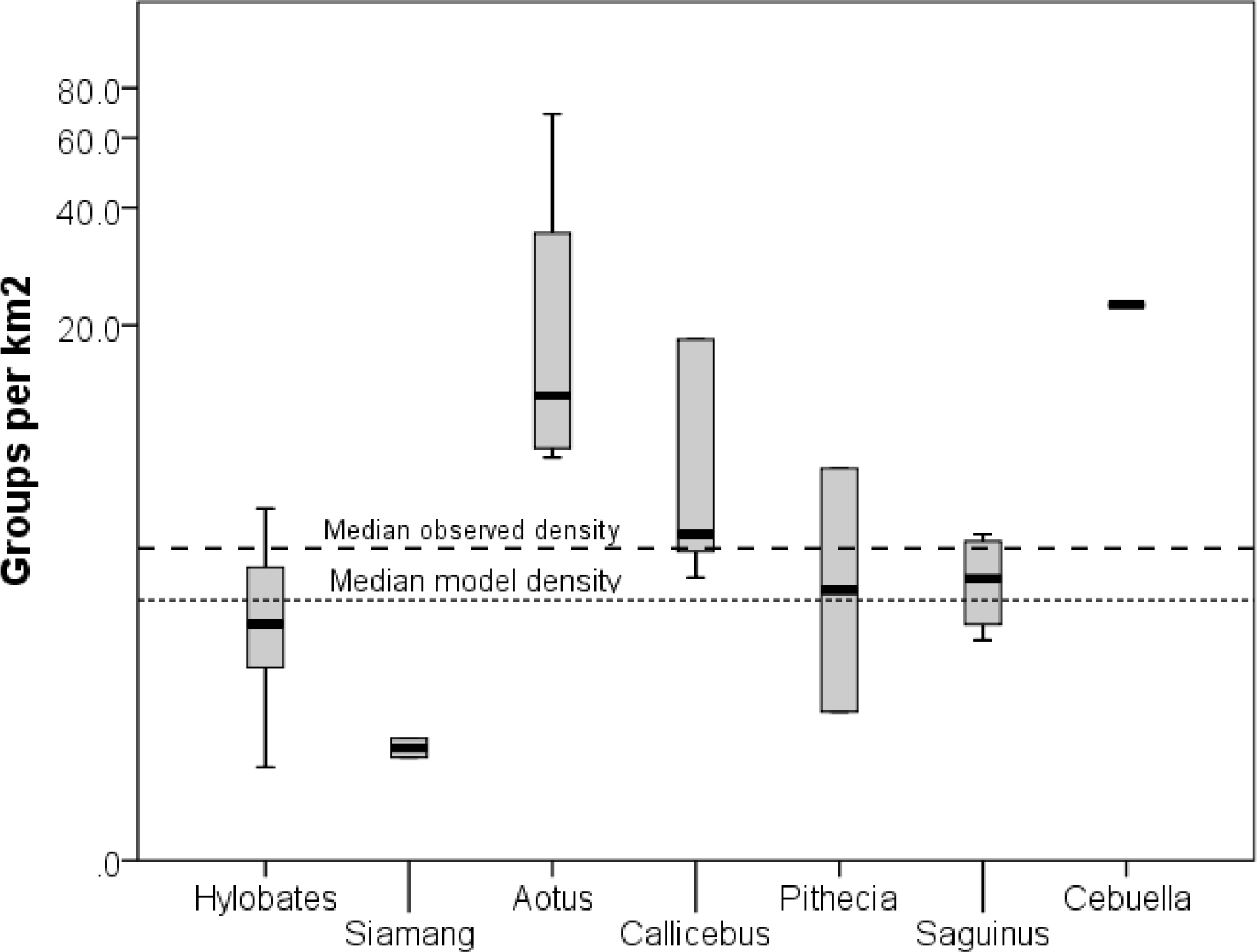
Median (±50% and 95% ranges) observed density of groups for monogamous genera. The dashed line indicates the median value for the observed data; the dotted line indicates the median value assumed in the model.

### Benefits of Roving

Fig. 4 plots the payoff ratio for a roving male (the number of offspring he could expect to sire compared to a monogamous male who sires one offspring in each interbirth interval). Data on interbirth intervals are not available for wild *Cercopithecus* populations or the prosimians, as well as a number of the monogamous genera (notably the hylobatids). Nonetheless, the sample is large enough to suggest a clear pattern: only *Colobus* and *Gorilla* would benefit by permanently attaching themselves to a group of females rather than opting for a promiscuous roving male strategy (i.e. their payoff ratios are not significantly above their observed female group size; one-sample *t*-tests, one-tailed in a positive direction: *Colobus* p=0.913, *Gorilla* p=0.992). For the other genera, all of whom are obligately monogamous, males would do significantly better by pursuing a roving male strategy than by being monogamous (one sample *t*-tests: p≤0.038).

**Fig. 4.**
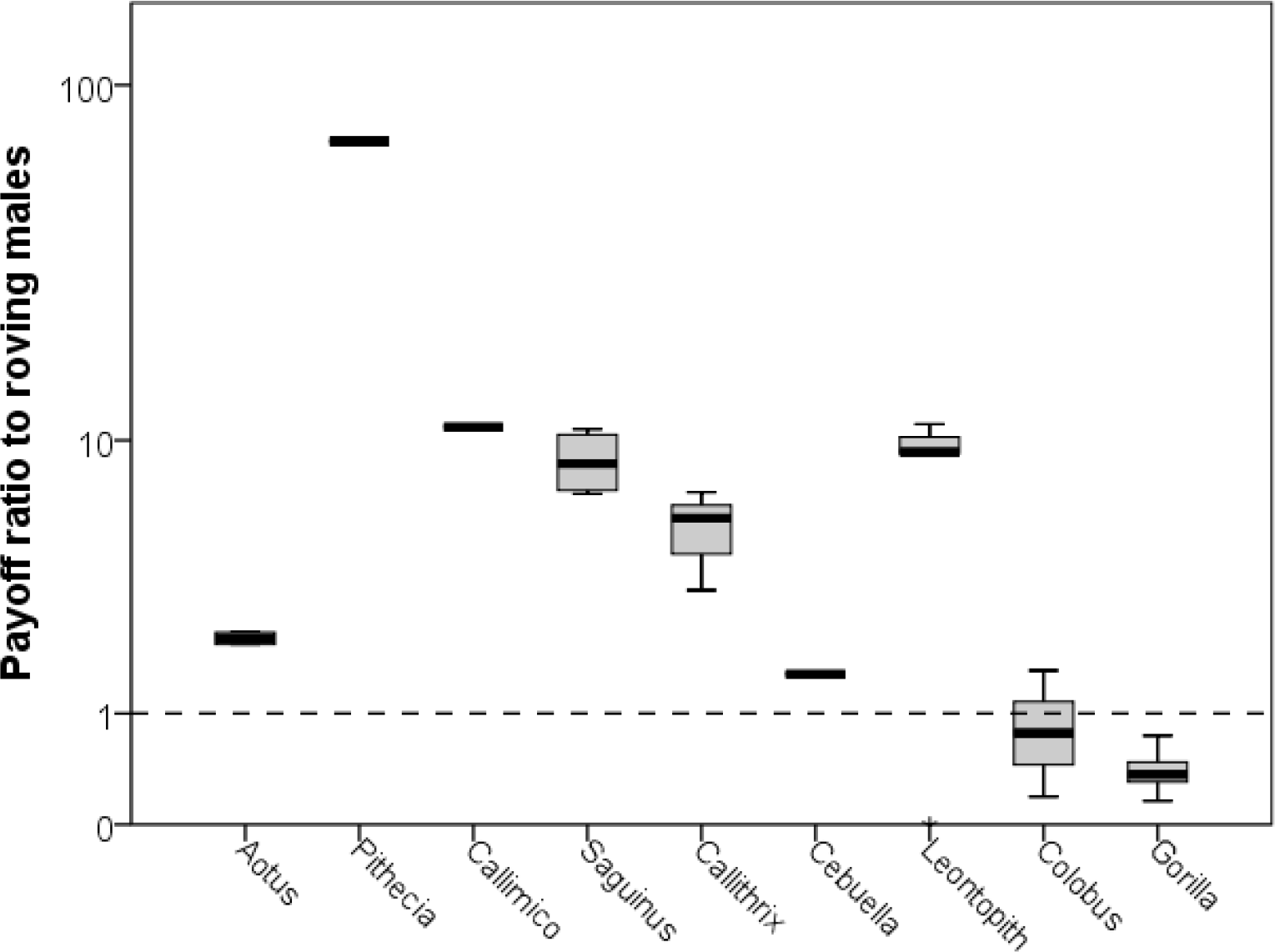
Median (±50% and 95% ranges) payoff ratio to promiscuous males (‘roving male strategy’) predicted by the male mating strategies model for the monogamous genera and two comparator polygynous genera (*Colobus* and *Gorilla*). The payoff ratio is the number of offspring sired in the average reproductive cycle (i.e. interbirth interval) by a promiscuous roving male divided by that expected for a social male (which equals female group size). The dashed line at a ratio of 1 indicates the point of equilibrium where the payoffs are equal and males should be ambivalent about which strategy to adopt. When the ratio lies above this line, males should opt for roving male polygyny; when it falls below, they should opt for being social (i.e. permanently attaching themselves to a group of females). When females live or forage alone, social males will be monogamous. The model correctly predicts that *Colobus* and *Gorilla* males should be social (and correctly predicts that orang-utan and chimpanzees males should be roving male polygamists).

## Discussion

All three tests suggest that the central assumption of the female dispersal hypothesis does not hold in the case of monogamous primates. With the exception of *Leontopithecus*, females of monogamous species do not occupy territories that are significantly larger than those required by females of polygamous primate species. Given the criterion that defines territoriality in primates, the males of most of these species could defend territories large enough to include the ranges of 5-6 solitary females, and could do at least as well as males from polygamous species that defend groups of females. In contrast to males from polygamous species, males from the obligately monogamous genera would all do better to adopt a form of roving male polygamy than to be monogamous.

In the case of the second and third test, the monogamous genera differ from baseline by a sufficient margin that it would require considerable error variance in respect of either the models or the data, or both, to nullify these results. In respect of Fig. 2, for example, estimates of day journey length or group density would have to be overestimated, or territory size underestimated, by a factor of 5-6 to bring the monogamous taxa down to baseline. This would be an implausible level of estimation error in any data. More importantly, *Colobus, Cercopithecus*, *Gorilla* and the prosimians provide us with confirmation that the models make the correct predictions: these genera behave exactly as they should do, preferring social polygyny to monogamy or a roving male strategy in the case of the anthropoids and a roving male strategy in preference to monogamy in the case of the prosimians. They provide us with an important element of ground-truthing for the models.

It would be difficult to argue with the conclusions of the first analysis, however: this explicitly tests the fundamental assumption that underlies the dispersed female hypothesis for monogamy – that females actually are overdispersed. One might argue that the second and third analyses are based on models, and models can be wrong or wrongly conceived. However, this claim would only be defendable if it could be shown that the models did not predict the general behaviour of primates especially well. In fact, both the Mitani-Rodman relationship and the male mating strategies model predict primate territorial and social behaviour extremely well, and particularly so in the case of Eq.[6]. If we want to question the validity the second model, we have to be able to explain away the fact that the model’s predictions fit the observed data extremely well, not just for apes but also for humans (Dunbar 2010) and for feral goats (Dunbar 1990).

This is not to say that solitary foraging by females is not an important feature of monogamy. When the reproductive advantages of doing so are strong enough, males attach themselves to female groups rather than opt to go roving. When females forage in groups, males become polygynous; when females forage alone, males are obliged to be monogamous. The issue is not that female ranges are so large that males cannot defend more than one female, but rather that some other selection pressure(s) constrains males into being monogamous whenever females forage on their own because they cannot afford to leave their female unguarded. If that pressure is low, males may prefer a roving strategy (as in the case of the nocturnal prosimians). Indeed, *Saguinus* males often do so, resulting in complex shifts in group structure and mating pattern over time (Goldizen 1988). However, they seemingly do so only when there are related follower males available to take over the group and provide the female with the rearing services she requires to maintain her high reproductive rate (Dunbar 1995a,b).

In primates, obligate monogamy appears to be a demographic and cognitive sink (Shultz et al. 2011). Once a species has opted for monogamy, it seems to be difficult for it to revert back to some form of polygyny, probably because the cognitive demands of monogamy (e.g. same-sex intolerance in both sexes) result in changes in brain organisation that cannot easily be undone. The opportunity cost that these males pay by being obligate monogamists is extremely high. By mating monogamously, they sire just one offspring in the average female reproductive cycle; by doing so they forego at least 4 offspring per reproductive cycle. In contrast, the nocturnal prosimians are willing to opt for roving male promiscuity even though the benefits of doing so are quite modest (on average, just 0.72 extra offspring per reproductive cycle: Fig. 2). This contrast reinforces the suggestion that the selection pressure against promiscuity is very high for monogamous anthropoids (and, by contrast, rather low for nocturnal prosimians).

The only two costs known to be high for anthropoids are predation (Shultz et al. 2004; Shultz & Finlayson 2010; see also Burnham et al. 2012) and infanticide by males (Opie et al. 2013). Such a claim need not apply to other mammals, where biological specificities may well result in female dispersion being the primary selection factor, as Lukas & Clutton-Brock (2013) have shown. The issue for primates is much more likely a consequence of the high risk they run from infanticide because of their greatly extended interbirth intervals, in turn a direct consequence of their unusually large brains and the fact that brain tissue can only be laid down at a slow, constant rate. The fact that the nocturnal prosimians do not opt for monogamy, despite high predation rates, tends to point to infanticide risk as the central problem. Uniquely, the nocturnal prosimians are able to duck this risk, partly because their smaller brains and more seasonal reproductive pattern mean shorter, less variable, interbirth intervals and partly because the females cache their vulnerable infants in nests rather than carrying them where they are protected from males.

## Acknowledgments

RD’s research is supported by a European Research Council Advanced grant. I thank Susanne Shultz for helpful suggestions.

## Appendix List of species included in the sample

List of species included in the sample.
|  | Species | Number of<br>populations sampled | Predominant<br>mating system |
| --- | --- | --- | --- |
| 484 | *Aotus azarai* | 3 | monogamy |
| 485 | *Aotus nigriceps* | 1 | monogamy |
| 486 | *Aotus trivirgatus* | 1 | monogamy |
| 487 | *Callicebus personatus* | 3 | monogamy |
| 488 | *Callicebus moloch* | 3 | monogamy |
| 489 | *Pithecia albicans* | 1 | monogamy |
| 490 | *Pithecia hirsuta* | 1 | monogamy |
| 491 | *Pithecia pithecia* | 3 | monogamy |
| 492 | *Callithrix torquatus* | 1 | monogamy |
| 493 | *Callithrix flaviceps* | 1 | variable (monogamy) |
| 494 | *Callithrix jacchus* | 3 | variable (monogamy) |
| 495 | *Callithrix kuhlii* | 1 | variable (monogamy) |
| 496 | *Callithrix pencillata* | 1 | variable (monogamy) |
| 497 | *Callimico aurita* | 2 | variable (monogamy) |
| 498 | *Callimico goeldii* | 2 | variable (monogamy) |
| 499 | *Cebuella pygmea* | 2 | variable (monogamy) |
| 500 | *Leontopithecus caissara* | 1 | variable (monogamy) |
| 501 | *Leontopithecus chrysomelas* | 2 | variable (monogamy) |
| 502 | *Leontopithecus chrysopygus* | 2 | variable (monogamy) |
| 503 | *Leontopithecus rosalia* | 2 | variable (monogamy) |
| 504 | *Mico argentatus* | 2 | variable (monogamy) |
| 505 | *Mico intermedius* | 1 | variable (monogamy) |
| 506 | *Saguinus fuscata* | 4 | variable (monogamy) |
| 507 | *Saguinus imperator* | 1 | variable (monogamy) |
| 508 | *Saguinus midas* | 1 | variable (monogamy) |
| 509 | *Saguinus mystax* | 2 | variable (monogamy) |
| 510 | *Saguinus tripartitus* | 1 | variable (monogamy) |
| 511 | *Hoolock hoolock* | 3 | monogamy |
| 512 | *Hylobates agilis* | 1 | monogamy |
| 513 | *Hylobates klossii* | 1 | monogamy |
| 514 | *Hylobates lar* | 4 | monogamy |
| 515 | *Hylobates moloch* | 1 | monogamy |
| 516 | *Hylobates muelleri* | 1 | monogamy |
| 517 | *Hylobates pileatus* | 1 | monogamy |
| 518 | *Nomascus concolor* | 1 | monogamy |
| 519 | *Nomascus leucogenys* | 1 | monogamy |
| 520 | *Symphalangus syndactylus* | 4 | monogamy |
| 521 | *Cercopithecus cephus* | 1 | monogamy |
| 522 | *Cercopithecus neglectus* | 1 | monogamy |
| 523 | *Cercopithecus ascanius* | 1 | unimale polygyny |
| 524 | *Cercopithecus diana* | 1 | unimale polygyny |
| 525 | *Cercopithecus l'hoesti* | 1 | unimale polygyny |
| 526 | *Cercopithecus mitis* | 3 | unimale polygyny |
| 527 | *Cercopithecus nictitans* | 2 | unimale polygyny |
| 528 | *Cercopithecus pogonias* | 1 | unimale polygyny |
| 529 | *Colobus angolensis* | 1 | unimale polygyny |
| 530 | *Colobus guereza* | 5 | unimale polygyny |
| 531 | *Colobus polykomos* | 2 | unimale polygyny |
| 532 | *Colobus satanas* | 2 | unimale polygyny |
| 533 | *Gorilla gorilla* | 3 | unimale polygyny |
| 534 | *Gorilla graueri* | 1 | unimale polygyny |
| 535 | *Gorilla beringei* | 1 | unimale polygyny |
| 536 | *Daubentonia* sp. |  | roving male polygyny |
| 537 | *Microcebus murinus* |  | roving male polygyny |
| 538 | *Perodictus potto* |  | roving male polygyny |
| 539 | *Loris lydekkerianus* |  | roving male polygyny |
| 540 | *Nycticebus coucang* |  | roving male polygyny |
| 541 | *Galagoides demidovii* |  | roving male polygyny |
| 542 | *Galagoides cocos* |  | roving male polygyny |
| 543 | *Galagoides moholi* |  | roving male polygyny |
| 544 | *Sciurocheirus gabonensis* |  | roving male polygyny |
| 545 | *Sciurocheirus alleni* |  | roving male polygyny |
| 546 | *Otolemur garnetti* |  | roving male polygyny |

